# Structured surveys of Australian native possum excreta predict Buruli ulcer occurrence in humans

**DOI:** 10.1101/2022.11.16.516821

**Authors:** Koen Vandelannoote, Andrew H. Buultjens, Jessica L. Porter, Anita Velink, John R. Wallace, Kim R. Blasdell, Michael Dunn, Victoria Boyd, Janet A. M. Fyfe, Ee Laine Tay, Paul D. R. Johnson, Saras Windecker, Nick Golding, Timothy P. Stinear

## Abstract

Buruli ulcer (BU) is a neglected tropical disease caused by infection of subcutaneous tissue with *Mycobacterium ulcerans*. BU is commonly reported across rural regions of Central and West Africa but has been increasing dramatically in temperate southeast Australia around the major metropolitan city of Melbourne. Previous research has shown that Australian native possums are reservoirs of *M. ulcerans* and that they shed the bacteria in their fecal material (excreta). Field surveys show that locales where possums harbor *M. ulcerans* overlap with human cases of BU, raising the possibility of using possum excreta surveys to predict the risk of disease occurrence in humans. We thus established a highly structured 12-month possum excreta surveillance program across an area of 350 km^2^ in the Mornington Peninsula area 70 km south of Melbourne, Australia. The primary objective of our study was to assess if *M. ulcerans* surveillance of possum excreta provided useful information for predicting future human BU case locations. Over two sampling campaigns in summer and winter, we collected 2282 possum excreta specimens of which 11% were PCR positive for *M. ulcerans*-specific DNA. Using the spatial scanning statistical tool *SatScan*, we observed non-random, co-correlated clustering of both *M. ulcerans* positive possum excreta and human BU cases. We next trained a statistical model with the Mornington Peninsula excreta survey data to predict the future likelihood of human BU cases occurring in the region. By observing where human BU cases subsequently occurred, we show that the excreta model performance was superior to a null model trained using the previous year’s human BU case incidence data (AUC 0.66 vs 0.55). We then used data unseen by the excreta-informed model from a new survey of 661 possum excreta specimens in Geelong, a geographically separate BU endemic area to the southwest of Melbourne, to prospectively predict the location of human BU cases in that region. As for the Mornington Peninsula, the excreta-based BU prediction model outperformed the null model (AUC 0.75 vs 0.50) and pinpointed specific locations in Geelong where interventions could be deployed to interrupt disease spread. This study highlights the *One Health* nature of BU by confirming a quantitative relationship between possum excreta shedding of *M. ulcerans* and humans developing BU. The excreta survey-informed modeling we have described will be a powerful tool for efficient targeting of public health responses to stop BU.

## INTRODUCTION

Buruli ulcer (BU) is an infection of subcutaneous tissues that can leave patients with disability and life-long deformity. The causative agent, *Mycobacterium ulcerans* is a slow-growing environmental bacterium that can infect humans after introduction through skin micro-trauma (1). While the exact environmental reservoir(s) and mode(s) of transmission of BU are unresolved, they do continue to be the subject of intense research. The disease has a highly focal geographical distribution and typically occurs around low-lying marshlands and riverine areas. BU occurs mostly in tropical and subtropical areas of West and Central Africa however smaller foci are recognized in South America, Southeast Asia and Australasia (2). In Australia, although small disease foci have been described in coastal regions of Queensland and the Northern Territory, the majority of the disease burden is found in the temperate state of Victoria, where several outbreaks have been recorded over the past three decades (3).

In Victoria, it has been established that the median incubation period is 4.8 months with peak BU transmission in humans occurring in late summer (4). Several cases of BU infections have also been reported in native wildlife and domestic mammal species including common ringtail (*Pseudocheirus peregrinus*), common brushtail (*Trichosurus vulpecula*), mountain brushtail possums (*Trichosurus cunninghami*) (5,6), koalas (*Phascolarctos cinereus*) (6), long footed potoroos (*Potorous longipes*), dogs (7), cats (8), horses (9) and alpacas (6). These naturally acquired BU infections in animals have occurred across the same geographical regions of Victoria where BU is known to be endemic for humans.

Several studies from Victoria suggest that BU is a zoonosis that first causes epizootic disease in the local native possum populations and then spills over to humans. Focused field surveys have revealed that possums excrete *M. ulcerans* DNA in their feces (excreta) in regions known to be endemic for humans while similar surveys outside endemic areas yielded negative results (10,11). While *M. ulcerans* DNA could be detected at low levels in a variety of other environmental samples in these studies, by far the highest concentrations of *M. ulcerans* were found in possum excreta (10). Subsequent capture and clinical assessment of free-ranging possums validated the findings of these excreta surveys by showing that subclinical *M. ulcerans* gut carriage in possums was common while a number of animals had laboratory-confirmed BU skin lesions and in some cases, advanced systemic disease (5).

Finally, whole genome sequencing and comparative genomic analysis has shown that *M. ulcerans* strains isolated from human and possum lesions are part of the same transmission cycles (10). These findings suggest that in Australia, BU is a One Health issue, with arboreal marsupial mammals representing an important environmental reservoir for *M. ulcerans*.

The present outbreak on the Mornington Peninsula in Victoria is the largest on record in Australia, with over 2,200 laboratory-confirmed cases diagnosed since 2010 (12). Concurrent with the Mornington Peninsula outbreak, has been the emergence in 2019 of a significant cluster of BU in a suburb of the major regional city of Geelong near the Bellarine Peninsula. Regions of the Bellarine Peninsula became a BU focus beginning in the late 1990s. The unprecedented increase in human BU cases since 2010 and the rapid expansion of BU endemic areas in Victoria, including incursions to within 5km of the Melbourne city centre (13) has highlighted how new strategies to control transmission are urgently required. An effective BU prevention and control program requires up-to-date information on the distribution of the disease and its incidence. However, the use of traditional epidemiological surveillance methods to track the emergence and movement of BU disease foci is severely limited by the long 5-month incubation period of the disease in humans (4), which complicates attributing disease acquisition to a particular event or location. Surveillance programs of a number of zoonotic pathogens like *Borrelia burgdorferi*, West Nile virus (14), and Rabies virus (15) are increasingly exploring the use of wildlife sentinels to monitor disease emergence and spread. Thus here, we explore the use of systematic screening surveys of possum excreta as an early warning surveillance system to monitor BU emergence in the Mornington Peninsula and Geelong region of Victoria. We show how the detection of *M. ulcerans* DNA in possum excreta was associated with the outbreak of BU disease in humans and we use statistical modeling to explore the potential of this approach as a public health tool to predict future BU emergence.

## METHODS

### STUDY SITE

The Mornington Peninsula is located 70km south of Melbourne’s city center and occupies a 750km^2^ area that separates Port Phillip Bay from Western Port Bay. The peninsula was one of the first areas in Victoria to be explored and settled by Europeans in the early 19^th^ Century. Since then, much of the native vegetation of the peninsula was cleared under pressure from urban development although fragmented pockets of remnant wild habitat have been preserved in the Mornington Peninsula National Park and are located southwest of the peninsula **(Figure S1).** The study site overlaps with the western tip of the Mornington Peninsula and is characterized by calcareous sandy soils that support a dense coastal scrubland. As the western tip of the peninsula is primarily a local tourist hotspot known for its affluent coastal resorts, a large proportion of the houses in the region serve as temporary tourist accommodation. Many of the residential properties in the study area are spacious holiday homes, set in fenced gardens planted with shrubs and trees, which represent ideal possum habitats.

To the west of the Mornington Peninsula, and on the opposite side of the Port Phillip Bay, lies Geelong at the eastern end of Corio Bay and the left bank of the Barwon River, approximately 65 km southwest of Melbourne. Geelong has an estimated urban population of 201,924 (as of June 2018) and is the second largest city in Victoria after Melbourne (16). Since 2019, BU has been considered endemic in the Geelong suburb of Belmont and surrounding areas, with local transmission suspected.

### ELECTRONIC DATA COLLECTION

Large-scale electronic data collection was organized using the “Build”, “Collect” and “Aggregate” tools of “Open Data Kit” (ODK), an open-source suite of tools that was designed to build data collection platforms (17). We used ODK Build to convert paper survey forms into an ODK compatible “electronic form”. The ODK Collect Android app was installed from the Google Play store onto five Android budget smartphones (hereafter referred to as survey phones). ODK Collect was then configured with the custom electronic form that contained all survey questions. During sampling surveyors worked their way through the prompts of the form and answered a wide range of question-and-answer types **(Figure S2).** The survey phones automatically sent finalized submissions over mobile data to an *ODK Aggregate* instance that was running in the cloud. Our instance of *ODK Aggregate* was hosted on the “Google Cloud Platform” cloud provider. All data collected by the survey phones was stored and managed on this platform. *ODK Aggregate* also automatically published all new database entries to *Google Sheets. We* used a custom script (18) to generate dynamic maps from collected data in this Google spreadsheet using the Google Maps API. This allowed a team-leader to monitor the progress of multiple teams surveying in real-time. After completing the survey all collected data was exported from the *ODK Aggregate* instance in either a tabular (csv) or a geographical (kml) format.

### STANDARDIZED ROADSIDE COLLECTION OF POSSUM EXCRETA

Samples of possum excreta were collected along the Mornington Peninsula Road network, which is mainly made up of low-traffic, single-track paved roads and unpaved gravel tracks. Samples were collected from the ground level along the fence line of residences on grassy strips and driveways along the road. To prevent re-sampling excreta from the same possum between adjacent points a sampling spacing interval of 200m was chosen which reflects the typical home-range (radius <100m) of these highly territorial animals (10). A 200m grid pattern was laid out with the help of a custom-built battery-operated distance tracker that incorporated an Arduino micro-controller, a GPS module, with an audio beep. When moving from sampling point A to sampling point B, the distance tracker would be reset at point A after which the device would measure 200m as the crow flies and report decreasing distance to point B by increasing the beeper’s intermittent beeping rate. ODK Collect was used to capture the location of the new sampling point using the survey phone’s GPS (accuracy <8m) after which the surveyor’s name was recorded. Surveyors then used the app to log a time point when they started looking for possum excreta. The search was restricted to a 50m radius around the sampling point and was terminated in case no excreta could be found after a pre-allotted time of 5min. Surveyors logged another time point when they discovered the first fecal pellet. Fecal material (excreta) from each sampling location was stored in separate sterile re-sealable plastic zipper bags that had been pre-labeled with a barcode. If possible, up to three excreta samples were sampled and, in case more were available, the freshest most intact excretas were selected. ODK Collect was then used to take a picture of the sampling site and scan the barcode on the zipper bag, both using the survey phone’s camera Following this, the collected excreta was used to distinguish and record the presumed species of possum based on distinctive morphological characteristics of the excreta from each species **(Figure S2).** Subsequently, completed forms were marked as finalized and uploaded to the cloud. In case no excreta material was found on a particular sampling location after 5min, ODK Collect forked to the end of the survey where the form likewise was marked as finalized and submitted. A video was produced and uploaded to YouTube to illustrate the steps described above (19). Samples were transported at 4°C to the laboratory where they were stored at - 20°C prior to further processing.

Several sampling missions were organized from 19 December 2018 to 14 March 2019 and are hereafter referred to as the “summer survey”. Later, between 28 May and 19 September 2019, we attempted to revisit most of the locations we sampled during summer, a period that is hereafter referred to as the “winter survey”. To facilitate the sampling effort during winter, the above-mentioned electronic distance trackers were reprogrammed with a predetermined grid of sampling points from the summer survey. Consequently, the trackers now helped surveyors relocate the nearest sampling point using the same auditory cue system. During the winter survey, we also tried to determine the usefulness of surveying at a higher resolution by sampling along a 50m grid pattern in three small regions. Apart from this, standardized ODK Collect-based roadside collection was performed identically. Another survey mission centered on the Geelong area was conducted in the early half of 2020 (16^th^ January through to the 28^th^ of April 2020) using the abovementioned surveying methodology with a 200m grid.

### DNA EXTRACTION, *M. ULCERANS* IS*2404* qPCR

For excreta samples collected from the Mornington Peninsula, microbial genomic DNA (gDNA) was extracted from possum excreta samples using the DNeasy PowerSoil HTP 96 Kit (Qiagen Cat# 12955-4) following the manufacturer’s protocols just prior to the addition of solution C4, whereupon DNA was subsequently purified from 200 μl of the lysate using the QIAsymphony DSP Virus/Pathogen extraction kit (Qiagen Cat# 937036) on the QIAsymphony automated platform. Extraction included two rounds of mechanical homogenization for 45 s at 1800□rpm on a FastPrep-96 instrument (MP Biomedicals). The extraction method for the possum excreta survey of the Geelong region followed a similar procedure but with some modifications as described elsewhere (20).

Real-time PCR assays targeting IS*2404* multiplexed with an internal positive control (IPC) (Life Technologies Cat# 4308323) were carried out as described before (21). Briefly, IS*2404* real-time PCR mixtures contained of 10.0□μl of 2x SensiFast Probe NO-ROX mix (BioLine Cat# BIO-86005), 3.2□μl of nuclease-free water, 0.8□μl each of 10□μM IS*2404* TF and IS*2404* TR primers, 0.8□μl of 5□μM IS*2404* TP probe, 2.0□μl TaqMan Exogenous IPC MIX, 0.4 μl TaqMan Exogenous IPC DNA, and 2.0□μl of DNA extract in a final reaction volume of 20□μl. Positive (*M. ulcerans* DNA) and negative controls (nuclease-free water) were included in each assay. Amplification and detection were performed with the Light Cycler 480 II (Roche) using the following program: 1 cycle of 95°C for 5 min, and 45 cycles of 95°C for 10 s and 60°C for 20 s.

### GEOGRAPHICAL DATA ACQUISITION AND SPATIAL ANALYSIS

The 2011 Victorian mesh block boundaries dataset and the 2011 Victorian mesh block census population counts dataset were downloaded from the Australian Bureau of Statistics (ABS) website (https://www.abs.gov.au/how-cite-abs-sources). Mesh blocks are relatively homogeneous statistical units and represent the smallest geographical unit for which publicly available census data are tabulated by the ABS. The mesh block digital boundaries dataset is based on Australia’s national coordinate reference system, the Geocentric Datum of Australia (GDA94). Spatial information was analyzed and edited in the geographic information system (GIS) software QGIS v.3.16.7 (22). The 2011 census population count spreadsheet was joined to the mesh block boundaries using the unique mesh block ID’s. The centroid of all polygon mesh blocks was determined and their latitude and longitude in GDA94 was calculated. The state-wide geometric dataset was then down sampled to a more manageable size (3238 mesh blocks) for subsequent spatial statistical analysis by reducing it to the ABS level 2 Statistical Areas (SA2) that encompasses the Mornington Peninsula (Point Nepean and Rosebud – McCrae). A second geographical dataset was prepared for the Geelong area by selecting the SA2 areas for that region (Belmont, Corio - Norlane, Geelong, Geelong West - Hamlyn Heights, Grovedale, Highton, Newcomb - Moolap, Newtown, and North Geelong - Bell Park).

All GPS positions of sampling points visited during the excreta surveys were added to this GIS and projected to GDA94. QGIS v.3.16.7 (22) was used to generate the figures of the excreta survey results and the geographical distribution of BU cases in the Mornington Peninsula.

### HUMAN BU CASES

The Victorian Department of Health (DH) made a de-identified database extract available of all BU patients that were notified in the state between 1 January 2019 and 31 December 2020. A human case of BU is defined here as a patient who presented with a clinical lesion suggestive of BU in which *M. ulcerans* DNA was detected laboratories using *IS2404* qPCR (21,23). Note that BU has been a ‘notifiable condition’ in Victoria since 2004 and as of 1 January, 2011 DH has been collecting enhanced BU surveillance data in a centralized notifiable disease database using custom collection forms (24). DH extracted the data used in this study from this database and then geocoded and de-identified it at an aggregated mesh block level. As mesh blocks typically encompass between 30 and 60 dwellings the extract of the notifiable disease database was effectively anonymized. Variables used in the analysis included: date of notification, date of first symptoms onset, “type” of contact with endemic area (resident, holiday resident, visitor), and mesh block of address of residence/holiday house/visit at the time of notification.

We selected two populations of notified BU patients suspected of having been infected with *M. ulcerans* in the Mornington Peninsula during an “exposure interval” that aligned with the organized possum excreta surveys. The reported onset of disease was used to infer this exposure interval based on the mean incubation period of BU in Victoria of 143 days (IQR 101–171) (4). We define the incubation period here as the time between exposure to an endemic region and symptom onset. Patients were suspected of having been infected with *M. ulcerans* in the Mornington Peninsula if they were either a resident of the peninsula or if they visited the area and had not reported recent (<12 months) contact with any other known BU endemic areas in the state. BU patients who, at the time of notification or prior to symptom onset, were residing in a house situated in the Mornington Peninsula are referred to here as “residents”. “Holiday residents” on the other hand were not domiciled in the Mornington Peninsula but stayed in a holiday house situated there. Finally, “visitors” are defined here as BU patients who had their exposure recorded outside of their place of residence while visiting the Mornington Peninsula during a short stay. A population of cases were also selected whose exposure interval overlapped with the survey of the Geelong region.

## ETHICS

Ethical approval for the use of de-identified mesh block-level BU case local information in this study was obtained from the Victorian Government Department of Health Human Ethics Committee under HREC/54166/DHHS-2019-179235(v3), “Spatial risk map of Buruli ulcer infection in Victoria”.

## STATISTICAL ANALYSIS

### BASIC STATISTICAL TESTING

Statistical analyses were performed using R v4.0.3 (http://www.R-project.org/). Comparison of excreta IS*2404* positivity between sampling season or possum species was done using Fisher’s exact test. Comparison of mean Ct (IS2404) values between sampling season or possum species was done with an unpaired t-test while assuming equal variance (checked with an F-test).

### SPATIAL SCAN STATISTICS

We used SaTScan v9.7 (25) to analyze surveillance data with discrete spatial scan statistics. SaTScan tackles this by progressively “scanning” a circular window of variable size across space while noting the number of observed and expected observations inside this window at each location. For each scanned circular window, a log likelihood ratio (LLR) statistic is calculated based on the number of observed and expected cases within and outside the circle and compared with the likelihood under the null hypothesis. We investigated Mornington Peninsula surveillance data both of (i) notified human BU disease and of (ii) epizootic spread of *M. ulcerans* in possum excreta. For each dataset, we accepted both primary and secondary clusters, if (i) their corresponding p-values were less than 0.005 and (ii) secondary clusters did not overlap geographically with previously reported clusters with a higher likelihood. P-values were based on 9999 Monte Carlo simulations for each dataset, as suggested by Kulldorff *et al*. (25).

We performed geographical surveillance of human BU disease by detecting spatial disease clusters and assessing their statistical significance. To achieve this, we applied the Poisson probability model to our mesh block level data of notified BU case counts, which arose from a background population at risk that was derived from the 2011 population census. We limited the maximum cluster size to 10% of the total population at risk (corresponding to 14,481 inhabitants) so defined areas of risk would remain within a manageable size for targeted BU prevention campaigns with limited resources. Under the null hypothesis, BU cases are homogeneously distributed over the Peninsula. Under the alternative hypothesis, there are geographical areas with higher rates of BU than would be expected if the risk of contracting BU was evenly distributed across the Peninsula.

We used the Bernoulli probability model to scan for spatial clusters with high rates of *IS2404* positivity in sampled possum excreta. The Bernoulli model is proper here as excreta *IS2404* positivity is a variable with two categories. The maximum cluster size was set to 50% of the population size. Under the null hypothesis, excreta *IS2404* positivity is homogeneously distributed over the surveyed region. Under the alternative hypothesis, there are clusters where the *IS2404* positivity rate is higher than in regions outside of these clusters.

### PREDICTIVE MODELING

To prospectively predict the occurrence of human BU cases, a statistical model was calibrated using excreta positivity and cases whose exposure period overlapped with the excreta survey periods. Specifically, cases were included if the exposure interval overlapped with the date that excretas were collected. Here the model was fitted separately for the summer and winter seasons with the objective to predict if a given mesh block will contain one or more human BU cases. The metric used to evaluate predictive performance was the area-under-the-curve (AUC) which is the degree of separability that describes how capable a model is at classifying mesh blocks where cases occur and those where cases don’t occur. AUC values range from 0 to 1, with AUC of 1 indicating a perfect record of classification while a value of 0.5 depicts a model that has no classification capacity. All scripts used in the statistical analytical pipeline have been made available in a GitHub Repository (26) and made use of the following R packages: flexclust (27), raster (28), readxl (29), sf (30), and tidyverse (31).

We built a custom statistical model to predict the probability of observing one or more human BU cases as a function of the distance-weighted prevalence of MU positivity in nearby excretas. That is, the probability of observing a BU case in each location is greater if nearby excretas are MU-positive. One approach to modelling the risk to humans would be to construct a geostatistical model of the prevalence of MU in excretas, i.e. a spatially continuous estimate of excreta positivity for all areas using e.g., model-based geostatistics (32). However, to use such a complex model for prediction of human cases to guide interventions would require re-estimation of the model and its many location-specific random-effect parameters after every trapping round. This would be burdensome and limit the application of the model in a public health setting. Instead, we construct a simpler model with two parameters that calculates the local prevalence around a focal location (e.g., the centroid of a small district) as the weighted mean of the excreta positivity in all samples, with those weights decaying with increasing distance from the focal location. The statistical model we use is as follows:

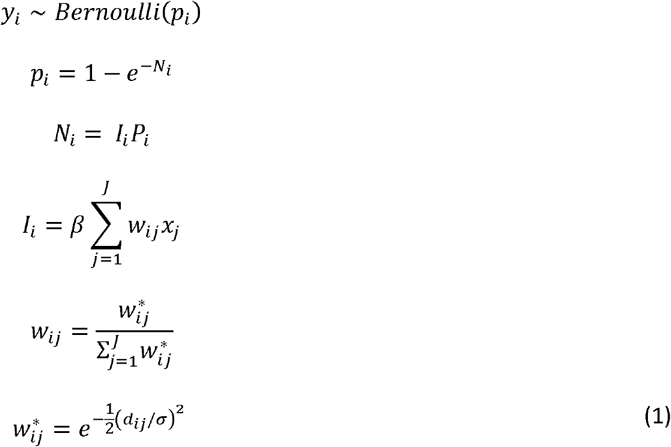

where *y_i_* is an indicator variable for whether a human BU case was detected at location *i* in the subsequent period, modelled as a single binomial sample with probability *p_i_*, calculated as the probability of observing one or more cases if case numbers in that location were Poisson-distributed with an expectation of *N_i_* cases; the product of the expected incidence *I_i_* and the population *P_i_* at each location. The incidence is modelled as the product of a positive-constrained scale parameter *β*, and the distance weighted average of the MU positivity *x_i_* (coded as a 1 for MU-positive or a 0 for MU-negative) across all excreta sampling locations *j* in *J*. The applied weight is a normalized Gaussian function of the Euclidean distance *d_ij_* between human case location *i* and excreta collection location *J*, with the range of the Gaussian function given by the positive-constrained parameter *σ*. The model is therefore parameterized by only two parameters: *β* which controls the absolute probability of observing a BU case in any given location, and *σ* which controls the distance at which excreta MU positivity is predictive of BU cases – if *σ* is small, only nearby excreta MU positivity is predictive of BU cases, whereas if *σ* is large, MU positivity in excreta collections further away are also informative. These two parameters are estimated by maximum likelihood using the Nelder-Mead optimization routine in R, with 5 random starts. These two estimated parameters can then be used to predict the probability of BU case occurrence *p_i_* in new places and sampling rounds, by combining them with a new excreta sampling dataset.

Note that this model could easily be modified to be fitted to, and therefore predict, the number *N_i_* of BU cases at a given location. However, given the comparatively low number of reported cases in each affected mesh block in our study region, the probability of presence of one or more cases is likely to be a more useful metric for prioritization of interventions. In addition, estimation of the conditional variance parameter of such a count model would add considerable complexity to the model fitting process and limit its utility for applied public health prioritisation.

### CROSS VALIDATION MODEL DEVELOPMENT

We first validated the predictive ability of the statistical model using a cross validation approach using the excreta survey data from Mornington Peninsula region. Here the mesh block dataset was split into three spatially contiguous regions so that spatial block cross validation could be performed **(Figure S3).** The model was fitted on each possible 2/3 of the data and then used to predict the probability of observing one or more BU cases in the remaining 1/3.

### PREDICTIONS ON UNSEEN DATA

The model was then fitted to the entire Mornington Peninsula dataset and used to make predictions to a previously unseen dataset of the Geelong excreta survey and human BU case data. The Geelong survey was conducted in early 2020 (16th January through to the 28th of April 2020), with there being just three mesh blocks that had cases with exposure intervals overlapping with the Gelong survey period. Geelong is a city of 200,000 inhabitants, 65 km southwest of Melbourne **(Figure S1)**

### ALTERNATIVE MODEL

Null models were established for the Mornington Peninsula and Geelong datasets that used the incidence of human BU cases in mesh blocks in the year preceding to predict the likelihood of a mesh block to contain a case. The null models were included as alternative models that were not based on any insights from the excreta surveys and represent the type of inference that could reasonably made to anticipate future case occurrence from epidemiological data alone. From a statistical perspective, the previous year’s incidence null models can be considered as a different option for out-of-sample intercept-only models for model performance comparison purposes.

### Ranking mesh blocks to inform BU transmission risk assessments

To provide a metric for real world application (*i.e*. pinpointing a region for potential public health interventions), the fraction of cases contained within the top percentages of predicted probability ranked mesh blocks were calculated. Here, the mesh blocks were ordered according to decreasing predicted class probability, with the fraction of total cases present in the top 5%, 10%, 20% and 50% of ranked mesh blocks recorded.

Due to the existence of many mesh block probability values for the null models having equal values (mostly zero), a randomization approach was employed to eliminate sorting artifacts. Here, a random seed was used 100 times to add a column of random integers (range 1-100) to the data frames containing model predictions. This data column was then used to initially sort the matrix prior to sorting on the predicted probability values. After 100 sorts on the randomised column followed by sorting on the probability values, the order of each mesh block was recorded and used to calculate an average order value. A final sorting operation was performed on the average order value to determine the mesh blocks that were in the top percentages.

## RESULTS

### POSSUM EXCRETA SAMPLE COLLECTION AND IS2404 QPCR TESTING

Standardized, grid-sectored roadside collection of possum excreta at approximately 200m intervals was performed in all freely accessible residential areas of the western tip of the Mornington Peninsula **(Figure S1).** A total of 2402 locations were visited during the two sampling seasons (summer: November 2018 - February 2019 and winter: May 2019 – August 2019). We encountered copious amounts of possum excreta during the surveys. In only 120 of all visited locations (5%) no excreta were found within the allotted 5-min survey time for each site. This observation indicates that the residential areas of the Mornington Peninsula support a large possum population.

During the surveys, excreta from common ringtail and common brushtail possums (hereafter referred to as ringtail and brushtail possums) were identified. Brushtail possum excreta were less frequent than ringtail excreta. *M. ulcerans* DNA was detected by IS*2404* qPCR in 310 of all 2282 (13.6%) excreta specimens **(Table 1**).

**Table 1:** Overview of the Mornington Peninsula excreta surveys across two seasons and *M. ulcerans* IS*2404* PCR screening results.

| Season | Possum species | No. of samples positive | No. of samples tested | Positivity rate | Sites with no excreta found |
| --- | --- | --- | --- | --- | --- |
| Summer |  | 111 | 987 | 11% | 58 |
|  | Ringtail possum | 110 | 947 | 13% |  |
|  | Brushtail possum | 1 | 40 | 3% |  |
| Winter |  | 199 | 1295 | 15% | 62 |
|  | Ringtail possum | 192 | 1237 | 16% |  |
|  | Brushtail possum | 7 | 58 | 12% |  |

### IMPACT OF SEASON ON M. ULCERANS PRESENCE IN POSSUM EXCRETA

In southeast Australia, the majority of BU transmission occurs in the summer months (4). We therefore tested the hypothesis that significantly more *M. ulcerans*-positive possum excreta material would be collected during summer. However, we observed no significant difference between excreta IS*2404* positivity and the sampling season (p = 0.2933, Fisher’s exact test). Additionally, no difference was found between the proportion of *M. ulcerans* IS*2404* positive excreta specimens and possum species (p=0.1014, Fisher’s exact test).

### CHANGES IN M. ULCERANS CONCENTRATION IN POSSUM EXCRETA OVER TIME AND SPACE

We used previously established IS*2404* qPCR standard curves to estimate the *M. ulcerans* load in possum excreta material. The Ct values for IS*2404* qPCR ranged from 25.6 – 40.0, corresponding to an estimated *M. ulcerans* excreta load of 24,000 – 5 mycobacterial genomes per fecal pellet (33). Interestingly, we noted that sampling season had a small – but statistically significant – impact on the fecal mycobacterial load (t(308)=-2.4, p=0.0171). On average, *M. ulcerans* positive excreta analyzed in summer had a Ct(IS*2404*) that was 0.87 lower than excreta collected in winter, a difference which corresponds with a 1.8 times higher mycobacterial load in summer excreta material. We observed no statistically significant difference between the fecal mycobacterial loads of the two possum species **(Figure 1**).

**Figure 1:**
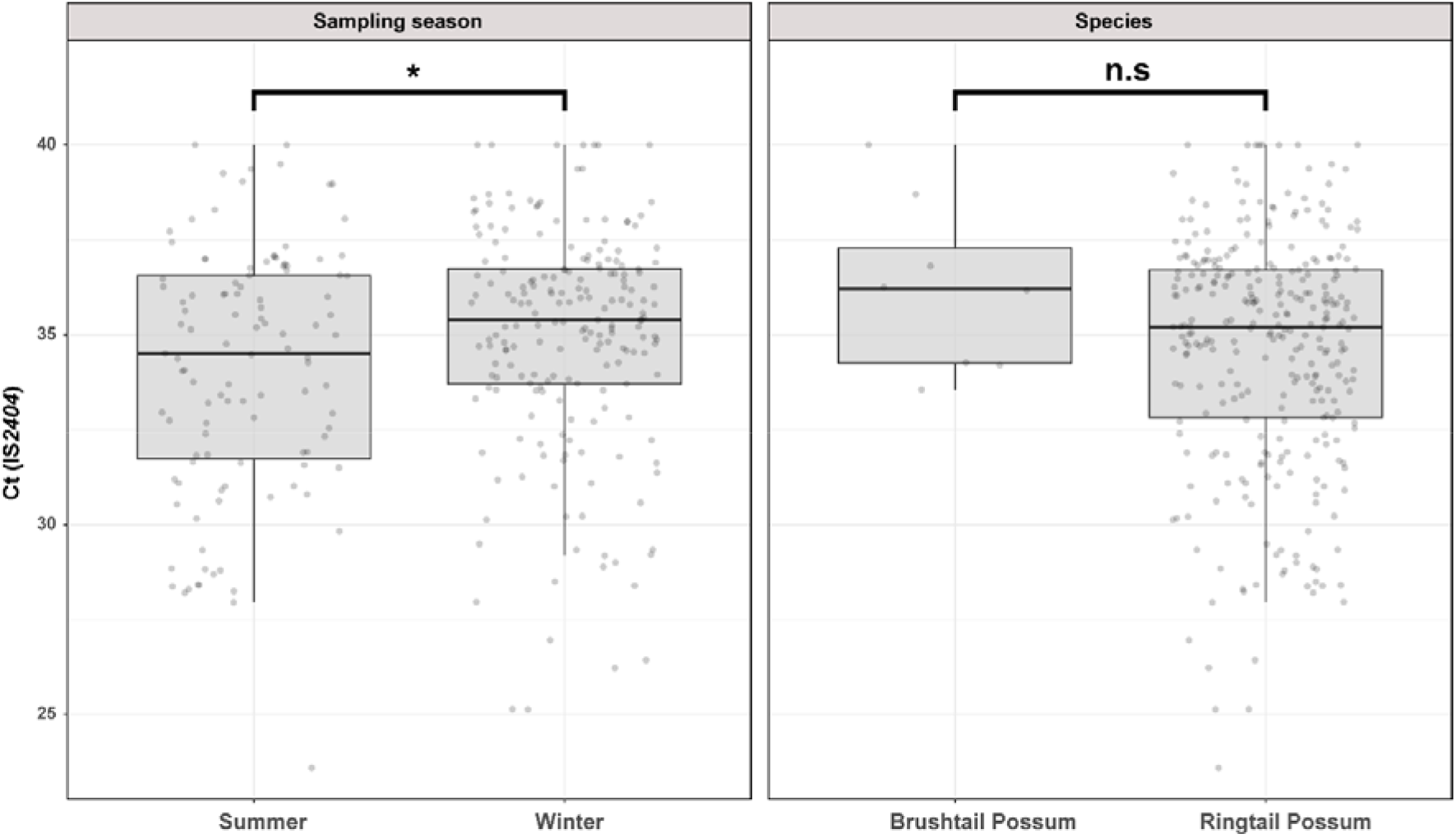
Overview of Mornington Peninsula M. ulcerans DNA concentrations in IS2404 positive excreta stratified by sampling season and possum species.

### SPATIAL DISTRIBUTION OF M. ULCERANS-POSITIVE POSSUM EXCRETA ACROSS THE MORNINGTON PENINSULA

The geographical distribution of *M. ulcerans* DNA in the Mornington Peninsula was investigated by mapping all GPS positions of sampling locations visited during the excreta surveys **(Figure 2 and 3).** Across the two sampling seasons, spatial scan statistics revealed three statistically significant geographical areas where IS2404 positive excreta clustered non-randomly **(Table 2).** Of note, the Sorrento SaTScan summer cluster encompassed an area where excreta with the highest *M. ulcerans* DNA concentrations of this study were also identified **(Figure 2).**

**Figure 2:**
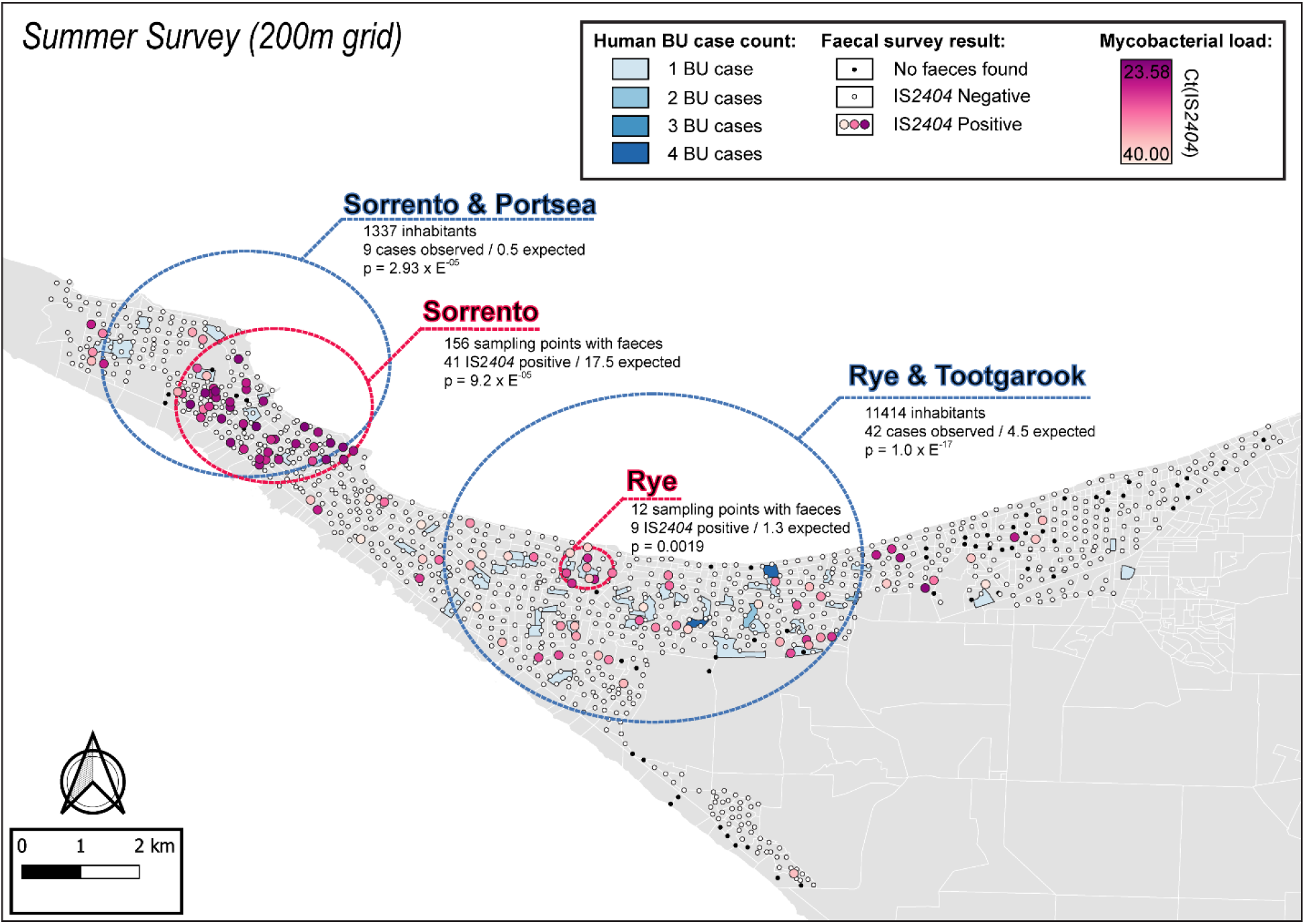
Geographical surveillance of M. ulcerans in the Mornington Peninsula during the southern hemisphere’s summer. The distribution of points where possum excreta was sampled along a 200m grid pattern is presented alongside with IS2404 molecular screening results. The pink to purple color gradient visualizes inferred mycobacterial loads in analyzed excreta as estimated from IS2404 cycle thresholds. The dashed red circles represent significant (p<0.005) non-random clustering of IS2404 positive possum excreta identified with spatial scan statistics. All BU patients notified to the DH with an inferred exposure time that overlapped with the excreta survey organized during summer are tabulated here by mesh block. A gradient is used to illustrate BU case counts per mesh block. The dashed blue circles represent geographical areas with higher rates of BU than would be expected if the risk of contracting BU was evenly distributed across the Peninsula.

**Figure 3:**
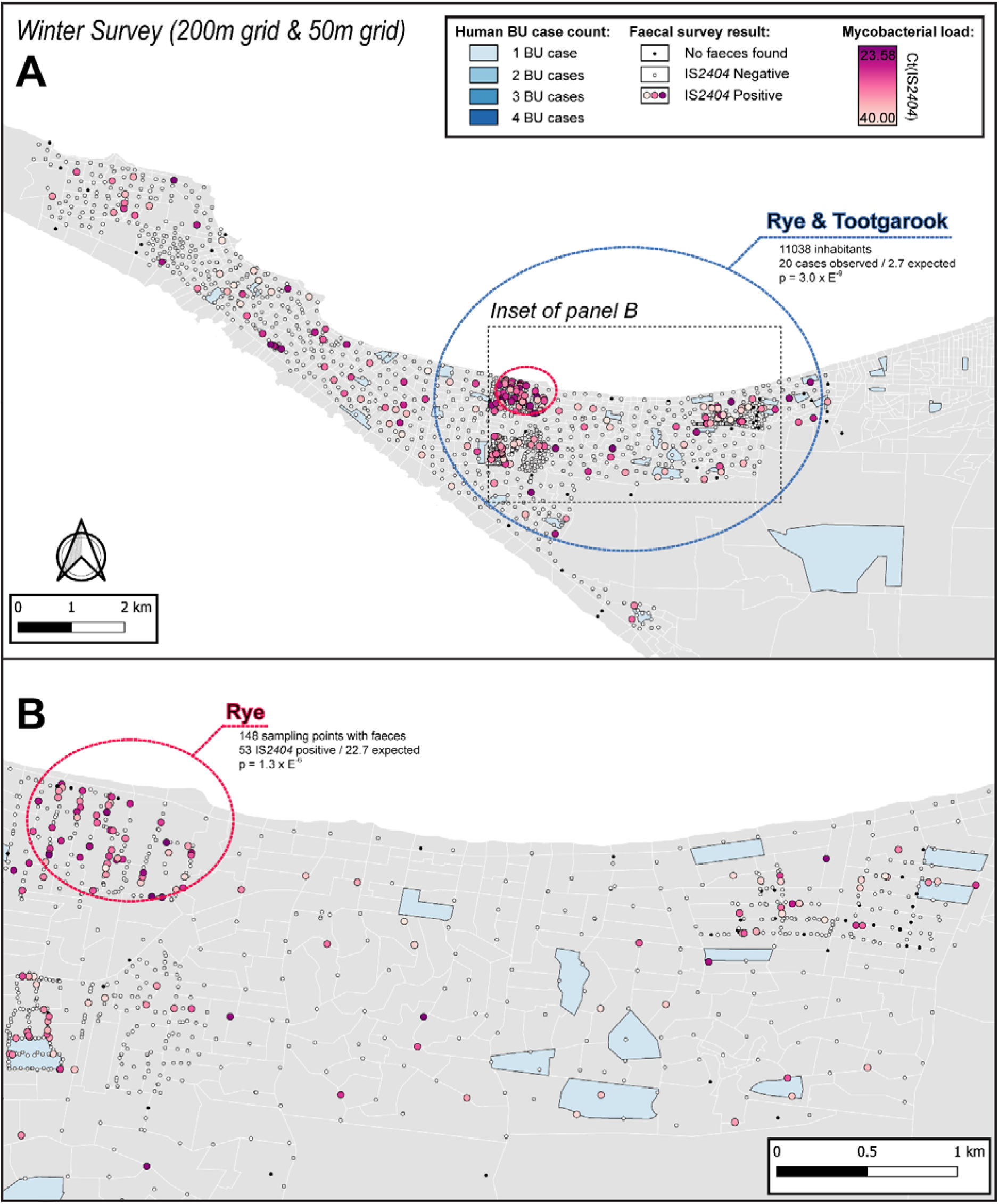
A&B: Geographical surveillance of M. ulcerans in the Mornington Peninsula during the southern hemisphere’s winter. The locations where standardized roadside collection of possum excreta was organized are illustrated alongside with IS2404 molecular screening results. The winter survey was performed along a 200m grid pattern although three limited regions were additionally sampled at higher resolution along a 50m grid (detailed in panel B). All BU patients notified to the DH with an inferred exposure time that overlapped with the excreta survey organized during winter are tabulated here by mesh block. The dashed circles represent geographical areas with higher rates of IS2404 positive possum excreta (red) or BU disease in humans (blue).

**Table 2:** Details of geographical clusters with high rates of *IS2404* positive possum excreta identified in SaTScan. LLR= Log Likelihood Ratio

| Survey | Approx. location | Cluster radius | # sampling locations | # IS2404 POS samples observed | # IS2404 POS samples expected | LLR | p-value |
| --- | --- | --- | --- | --- | --- | --- | --- |
| Summer | Sorrento | 1.7 km | 156 | 41 | 17.5 | 17.052 | 9.20E-05 |
| Summer | Rye | 0.4 km | 13 | 9 | 1.3 | 13.583 | 1.90E-03 |
| Winter | Rye | 0.6 km | 148 | 53 | 22.7 | 21.800 | 1.27E-06 |

### M. ULCERANS-POSITIVE POSSUM EXCRETA TO PREDICT OCCURRENCE OF BU IN HUMANS

A major aim of this research was to try and use the possum excreta survey data to predict the risk of BU transmission to humans. Our approach was to prospectively compare the non-random clusters of *M. ulcerans*-positive possum excreta samples described above with any non-random clusters of human BU cases that were likely acquired during the summer and winter sampling seasons. To do this, a de-identified DH database extract with enhanced surveillance data was used to select BU patients infected with *M. ulcerans* in the Mornington Peninsula during an exposure interval that aligned with the summer and winter excreta possum surveys **(Figure S4).** As a result, two populations of BU patients were identified that were highly likely to have been infected in the Mornington Peninsula during the summer (n=62) and winter (n=35) excreta surveys. On the assumption that residents/visitors/holiday residents were infected near their residences/holiday houses, the address of the property (geocoded to the 2011 mesh block level) was used in all spatial analyses. Additionally, the population-at-risk from which BU cases arose was represented by the Mornington Peninsula’s 144,817 inhabitants as recorded in the 2011 census.

Spatial scan statistics applied to these mesh block data for these BU cases revealed three statistically significant clusters across the two sampling seasons where human BU cases aggregated non-randomly. Across the two sampling seasons, there was a significant spatial correlation between the three clusters of human BU disease and the three clusters with high occurrence of *M. ulcerans* positive possum excreta **(Table 3, Figures 2 & 3).**

**Table 3:** Details of geographical clusters with high BU incidence identified in SaTScan. MB= mesh block, LLR= Log Likelihood Ratio

| Survey | Approx. location | Cluster radius | # MB centroids | Cluster census population | BU cases observed | BU cases expected | LLR | p-value |
| --- | --- | --- | --- | --- | --- | --- | --- | --- |
| Summer | Sorrento & Portsea | 2.4 km | 140 | 1337 | 9 | 0.5 | 17.749 | 2.94E-05 |
| Summer | Rye & Tootgarook | 3.6 km | 408 | 11414 | 42 | 4.5 | 75.087 | 1.00E-17 |
| Winter | Rye & Tootgarook | 3.6 km | 383 | 11038 | 20 | 2.7 | 28.770 | 3.05E-09 |

The statistical model developed to prospectively predict the probability of a mesh block containing a case demonstrated a superior ability under spatial-block cross-validation **(Figure S3)** to rank mesh blocks according to whether they contained a case or not during the exposure interval – the Receiver-Operating-Characteristic (ROC) area-under-the-curve (AUC) value - with a mean AUC of 0.66. This is compared to a mean AUC of 0.56 for a null model based on the preceding year’s human BU case incidence data for the Mornington Peninsula **(Figure 4A,B).** When fitted to the full Mornington Peninsula dataset, the model parameters were estimated as: *β* = 0.014 (95% confidence interval 0.01-0.02) and *σ* = 1.06 (0.92-1.21).

**Figure 4.**
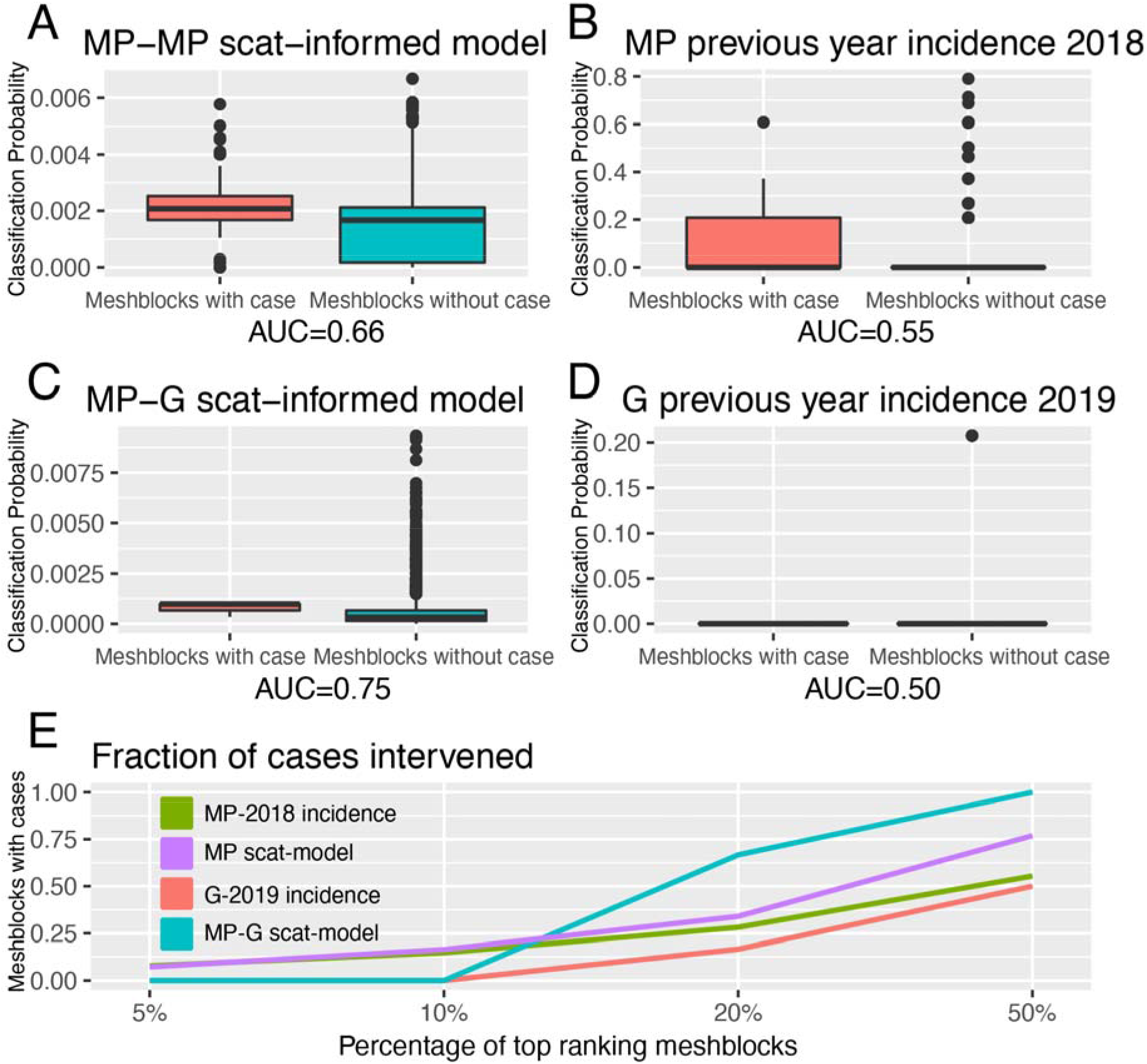
Statistical modeling approaches to prospectively predict the likelihood that a mesh block will contain a BU case. Mornington Peninsula and Geelong have been abbreviated as (MP) and (G), respectively. All paired boxplots show the predicted probabilities for mesh blocks with a case (red) and mesh blocks without a case (blue) and AUC value below each graph. A: Paired boxplot of the Mornington Peninsula excreta-informed model. B: Paired boxplot of the Mornington Peninsula excreta-informed model when predicting on the Geelong data. C: Paired boxplot of the Mornington Peninsula previous year’s incidence (2018) null model. D: Paired boxplot of the Geelong previous year’s incidence (2019) null model. E: Ranked performance of all predictive models. Ranking cutoff intervals included the top 5, 10, 20 and 50% of mesh blocks, ordered by their declining predicted class probabilities.

For the full out-of sample validation test, the model developed on the Mornington Peninsula data was validated against possum excreta survey data of 1,128 sites and 661 excreta specimens collected during 2020 in the Geelong region **(Figure S5),** and BU cases in the same region. Three confirmed human BU cases were reported from the Geelong region with a transmission interval that overlapped the sampling period **(Figures S5 and S6).** The model achieved an AUC of 0.75 **(Figure 4C),** compared with a null-model based on the previous year BU case incidence with an AUC score of 0.50, underscoring the predictive ability of the excreta-informed model **(Figure 4D).**

*M. ulcerans* DNA was detected by IS2404 qPCR in 21 of the 661 (3.2%) excreta specimens **(Table 4).** As was observed with the Mornington Peninsula survey, Common Ringtail possum excreta were more frequently found than Common Brushtail possum excreta but the proportion of IS2404 qPCR was not significantly different **(Table 4).**

**Table 4:** Geelong excreta survey results summary.

| Dates | Possum species | No. of samples positive | No. of samples tested | Positivity rate | Sites with no excreta found |
| --- | --- | --- | --- | --- | --- |
| January-April 2020 |  | 21 | 661 | 3% | 467 |
|  | Ringtail possum | 19 | 577 | 3% |  |
|  | Brushtail possum | 2 | 84 | 2% |  |

In addition to the AUC, we also calculated for all models the fraction of total mesh blocks with cases that were detected when targeting the top 5, 10, 20 and 50% of mesh blocks, ranked by the predicted probability that a mesh block will contain at least one BU case **(Figure 4E).** The excreta-informed models deployed in both the Mornington Peninsula and Geelong areas demonstrated a greater ability to classify case-containing mesh blocks into the 5, 10, 20 and 50% of top-ranking mesh blocks than that of the null-models **(Figure 4E).** Targeting of the top 20% of total mesh blocks for these two regions represents a substantial reduction in the overall number of mesh blocks at 368/1840 and 365/1827 for the Mornington Peninsula and Geelong models, respectively. Given the tradeoff between narrowing the geographic search area and maintaining sufficient sensitivity to detect mesh blocks where cases might occur, we found that the selection of the top 20% of excreta-informed model probability-ranked mesh blocks was a good compromise.

The geographical distribution of the top 20% of probability-ranked mesh blocks obtained with the excreta-informed models also had obvious spatial clustering compared with the null-models, which had excretatered mesh blocks **(Figure 5A-D).** The non-random mesh block probability density of the excreta-informed models suggests these data can be used to inform rational, targeted and thus more cost-effective deployment of any interventions compared with reliance on human case data alone.

**Figure 5:**
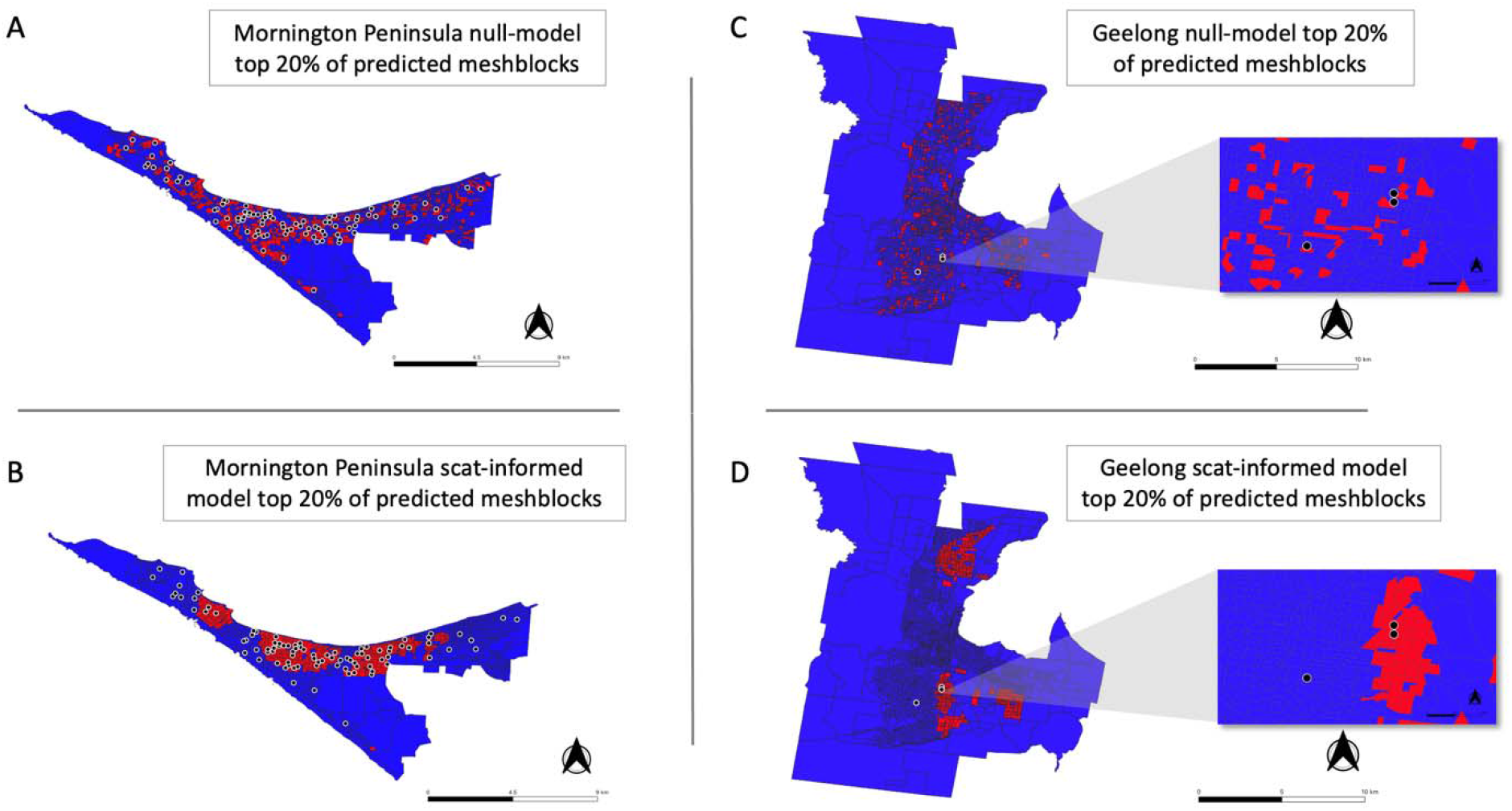
Geographical distribution of the top 20% of probability-ranked mesh blocks for all excreta-informed and null models. Red indicates mesh blocks in the top-20% of probability-ranked results while blue indicates mesh blocks in the bottom-80%. Black circles with white borders indicate location of confirmed BU cases that occurred within the transmission window following the excreta sampling. A: Mornington Peninsula null-model top-20% mesh block predictions. B: Mornington Peninsula excreta-informed model top-20% mesh block predictions. C: Geelong null-model top-20% mesh block predictions. D: Geelong excreta-informed model top-20% mesh block predictions. Insets show zoom-in of mesh blocks where BU cases occurred.

## DISCUSSION

The rapid expansion of BU endemic areas in southeastern Australia has highlighted the need for new strategies to understand and control disease spread. The long and variable incubation period of BU in humans has challenged traditional epidemiological surveillance approaches in tracking the emergence and movement of BU disease foci (4,34). Building on the findings of other zoonotic pathogen surveillance programs (14,15), here we explored the potential role arboreal marsupial mammals as wildlife sentinels to monitor BU emergence and spread and to help understand the role of these animals in the transmission of BU. Our primary goal was to determine whether *M. ulcerans* surveillance of possum excreta could act as an early warning system capable of predicting future human BU case locations. As a first task, we established a possum excreta surveillance program that monitored *M. ulcerans* in the environment of the Mornington Peninsula. This allowed us to determine the extent of epizootic activity during consecutive summer and winter seasons.

We identified a significant spatial correlation between clusters of *M. ulcerans* positive possum excreta and clusters of confirmed human BU cases likely infected with *M. ulcerans* in the same region during an exposure interval that aligned with the excreta possum surveys **(Figure 2).** While the overlap between the two cluster types was not perfect, it is important to highlight that the *SatScan* clusters detected represent the general area of a cluster and the circles are only approximate boundaries. Importantly however, the patterns and overlap we observed aligned with previous assessments of the positive association between *M. ulcerans* in possums and human BU cases (10,11,20) and thus very strongly implicate Australian native possums as key environmental reservoirs of *M. ulcerans*, involved in a transmission cycle with humans. This association between possums, human and BU flags this disease as a One Health issue. It also suggests that a surveillance program that monitors *M. ulcerans* DNA in possum excreta could alert public health authorities to increased human BU risks, which would allow prevention, and control programs to be implemented before human BU cases occur.

To further explore the potential of possum excreta surveys to predict the risk of BU cases occurring in humans in particular areas, we built a custom statistical model and compared its performance to null models built from the previous year’s human BU case incidence. From a public health perspective, the null models can be considered as a conventional approach to determine where cases might appear in the future. The development of a excreta-informed model for the Mornington Peninsula data revealed that it had greater predictive capacity than the Mornington Peninsula null model, in that it could more accurately predict areas (mesh blocks) with cases than the null model. We extended the Mornington Peninsula excreta-informed model to make predictions upon a previously unseen excreta survey dataset in the Geelong region. As for predictions made on the Mornington Peninsula, the predictions made with the Geelong excreta data had a greater ability to correctly predict human BU case-containing mesh blocks than was observed with a null model that used the previous year’s BU incidence.

This increased performance of the excreta-informed models might be explained by several factors. Firstly, the long incubation period makes it difficult to establish both when and where a person may have been infected with *M. ulcerans*, so the spatial information used by the null models will likely contain more ‘noise’ than the excreta-informed models. Possums, however, have a limited range, usually less than 100 m, and so their excreta is a spatially trustworthy analyte, providing a more accurate picture of pathogen distribution in the environment (10,35). Secondly, human BU cases are not detected in areas where possums do not harbour *M. ulcerans* (10,11,20). Therefore, possums are playing a substantially more contributive role than humans to BU transmission cycles, helping to explain why the excreta-informed model out-performs human BU case incidence-based models. The ability of the excreta-informed model trained with data from the Mornington Peninsula to predict human case occurrence in a distinct area (Geelong) strongly reinforces the link between possums harbouring *M. ulcerans* and human BU cases across different geographic areas in southeast Australia.

The ranking evaluation of the predictive models is another strength of the modeling approach, as it reports the number of mesh blocks where cases potentially could have been prevented (or better managed) if these top-ranking mesh blocks were targeted by an effective public health intervention. We envisage a scenario in which such geographical risk assessments could form the basis of public health messaging programs that target areas where disease transmission is predicted as most likely to occur. In this way, for instance, frontline general practice clinicians could be advised of the elevated local BU transmission risk, potentially leading to earlier patient diagnosis and improved clinical outcomes for the cases that do emerge. In addition, targeted messaging based on predictive modeling may also encourage preventative behaviors among local communities that could lessen the chances of transmission and reduce the number of emerging cases. For example, targeted messaging promoting behaviors that mitigate known BU risk factors (eg mosquito bites) or promote known protective behaviours (use of insect repellent) could reduce disease incidence (20,36). Other interventions could reduce the abundance of potential vectors that might be transmitting *M. ulcerans* from possums to people, or perhaps seek to control the infection in possums with novel therapeutic interventions.

Possum excreta faecal surveys are practical and cost effective because excreta from Australian native possums in urban and semi-urban areas is highly abundant, easily recognized and easily accessed. It is an ideal environmental analyte. Roadside collection of possum excreta and subsequent molecular screening is also relatively straightforward and processing epizootiological data does not require informed consent or access to medical records. Furthermore, as multiple years may elapse between the occurrences of BU in a particular region, continuous excreta surveillance might detect trends in the distribution and epidemiology of BU in a region and allow the effectiveness of any BU control measures; measures such as identifying and treating *M. ulcerans*-infected possum populations.

We explored the impact of sampling at a higher density (50m intervals instead of 200m) **(Figure 3A).** The higher density sampling provided by 50m intervals increased resolution for the faecal mapping, but it didn’t materially change the pattern of *M. ulcerans* positive faecal samples detected at 200m sampling grids. We propose that 200m sampling grids provide a pragmatic balance between survey sensitivity and survey time/costs.

The streamlined workflow, the custom-built distance trackers, and the use of Android mobile phones equipped with an electronic data collection solution simplified the fieldwork to such an extent that new field workers could be trained within a day. Moreover, incorporating ODK in our workflow allowed us to organize our mobile excreta surveys in a cost-effective manner as open-source software suite was hosted free-of-charge by a cloud provider. Furthermore, ODK Collect proved a powerful tool for ensuring high-quality data collection as it only uploaded completed submissions, as the app automatically validated survey responses at the point of data collection by using entry constraints, error checks and form logic. This significantly reduced data errors and data loss that would be much more common in paper surveys at this scale.

In this study, we have confirmed that Buruli ulcer in southeast Australia is a zoonotic infection involving Australian native possums in a transmission cycle. The means by which the pathogen is spread between possums and humans is under active investigation, but mosquitoes are likely vectors (37,38). We are also investigating the natural history of *M. ulcerans* infection in possums, to better understand the apparent susceptibility of these animals to mycobacteria. The detection of *M. ulcerans* DNA in possum excreta is associated with the occurrence of BU disease in humans and we have explored how these surveillance data can be used to predict future BU case emergence. A future surveillance program should collect, analyze, and interpret epidemiological, clinical, and epizootiological data on BU. Questions/issues to address about such a program include breadth of areas to survey, frequency of specimen collection, and availability of the resources/trained personnel required to establish and maintain a program. Environmental surveillance should identify epizootics as quickly as possible so that steps can be taken to control disease spread. We have found possum excreta surveillance of *M. ulcerans* DNA can identify the spread of BU epizootics and providing public health authorities with sufficient warning to implement control measures before human cases occur.

## ACKNOWLEDGMENTS

We are grateful to Gabrielle Stinear, Andrew Walker, Brianna Behrsin, Zoe Winkle, Zoe James, Simone Clayton, Kerri Howell and Jake A Linke for valued assistance during fieldwork.

## COMPETING INTERESTS

The authors have no competing interests to declare.

---

**Figure S1:**
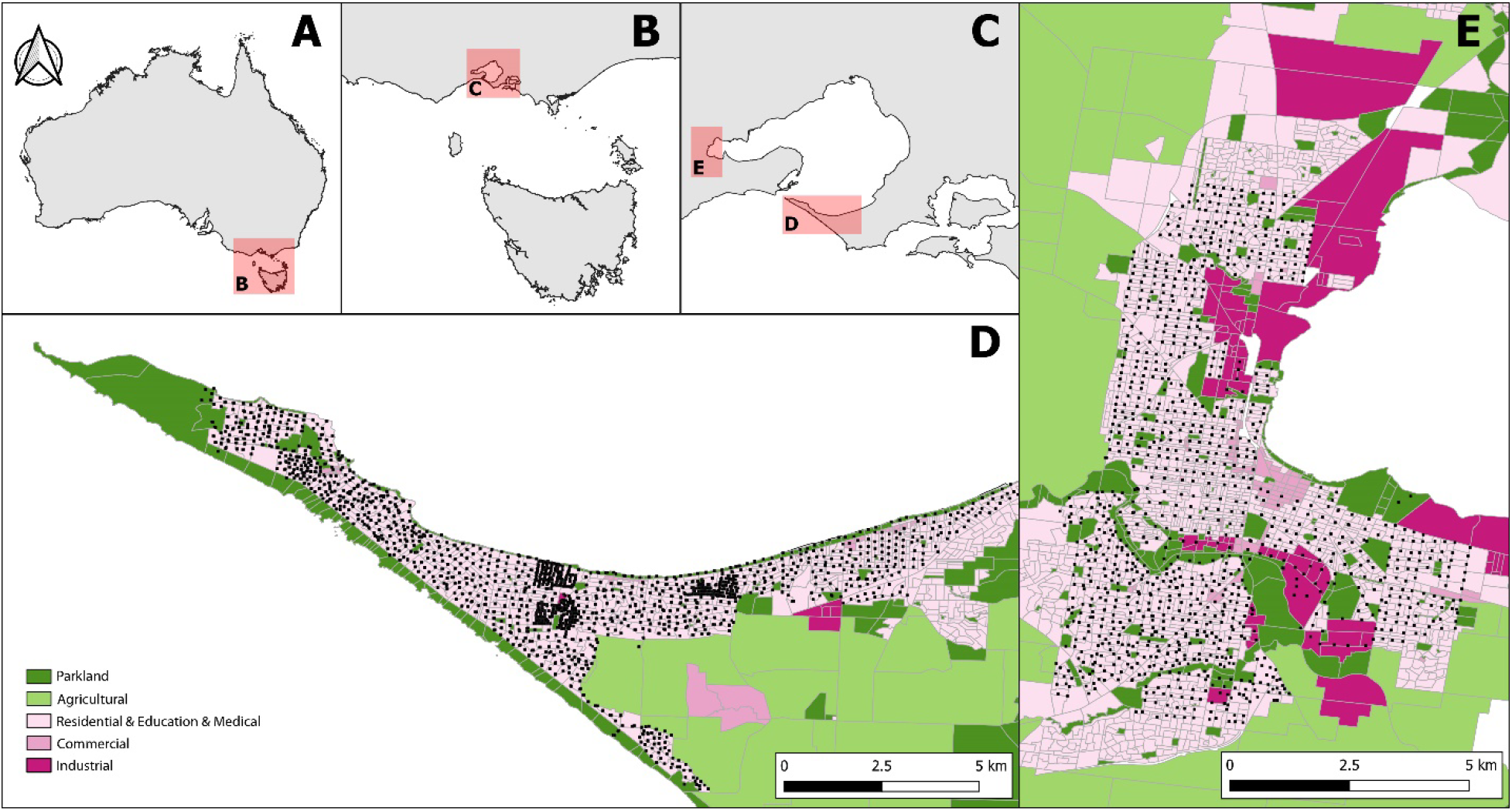
Geography of the Mornington Peninsula and Geelong. Panel A, B, C: zoomed inset maps at successively smaller scales are used to establish the location of the western tip of the Mornington Peninsula (Panel D) and Geelong (Panel E) in Australia. Panel D, E: land use categories of 2011 census mesh blocks as detailed in the legend. The GPS coordinates of all sample points visited during the study are rendered as black squares.

**Figure S2:**
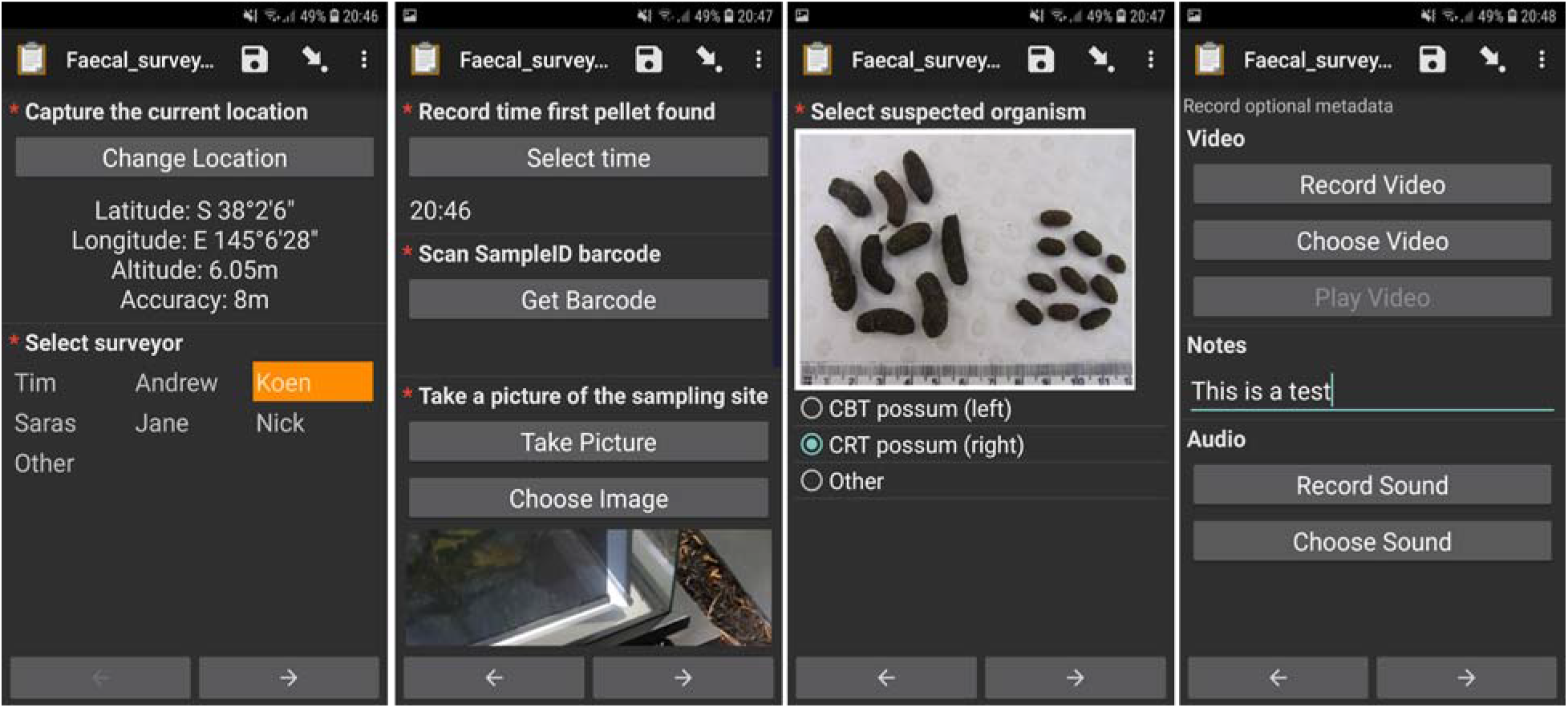
A set of screenshots demonstrating electronic data collection on the ODK Collect Android app running on a survey phone.

**Figure S3:**
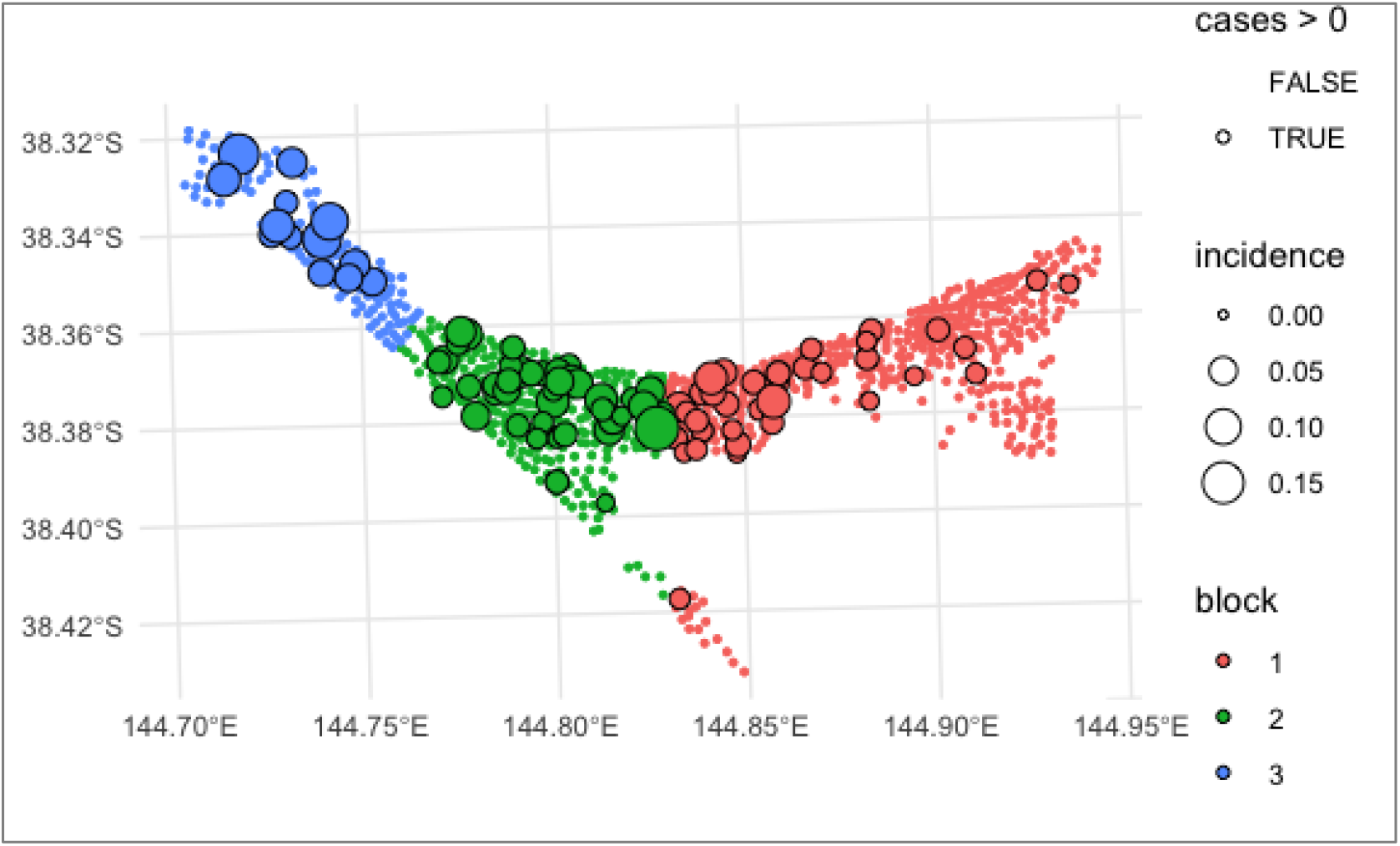
The three spatially contiguous blocks of the Mornington Peninsula mesh blocks that were devised for cross validation model development.

**Figure S4:**
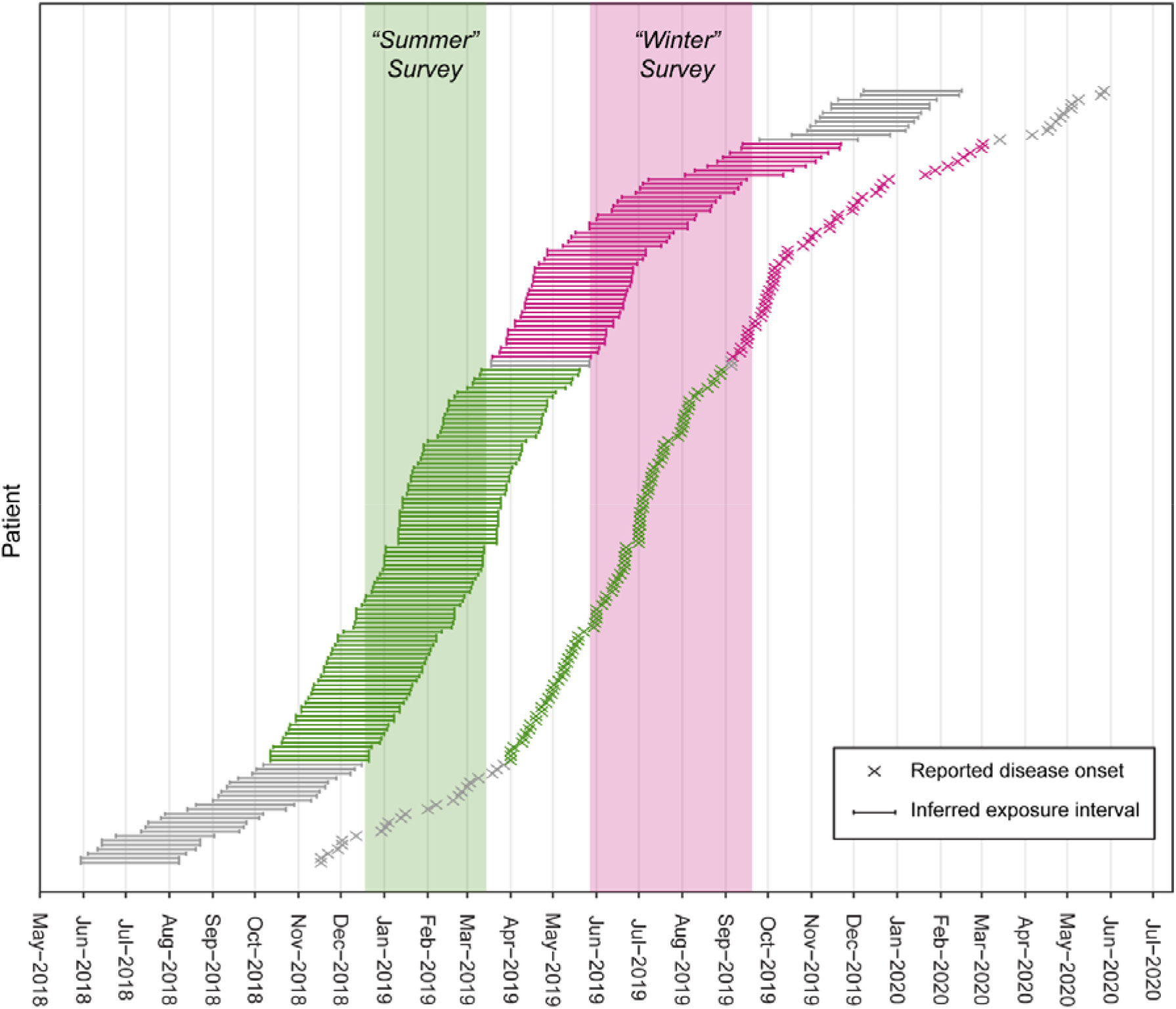
Selection of two populations of BU patients with exposure intervals that aligned with the excreta possum surveys organized during the southern hemispheres’ summer and winter. The two populations comprise BU patients notified to the Victorian DH who resided in or visited the Mornington Peninsula and had not reported recent (<12 months) contact with any other known BU endemic areas in the state. The reported onset of disease was used to infer the exposure interval during which patients were likely infected based on the mean incubation period of BU in Victoria of 143 days (IQR 101–171) (4).

**Figure S5:**
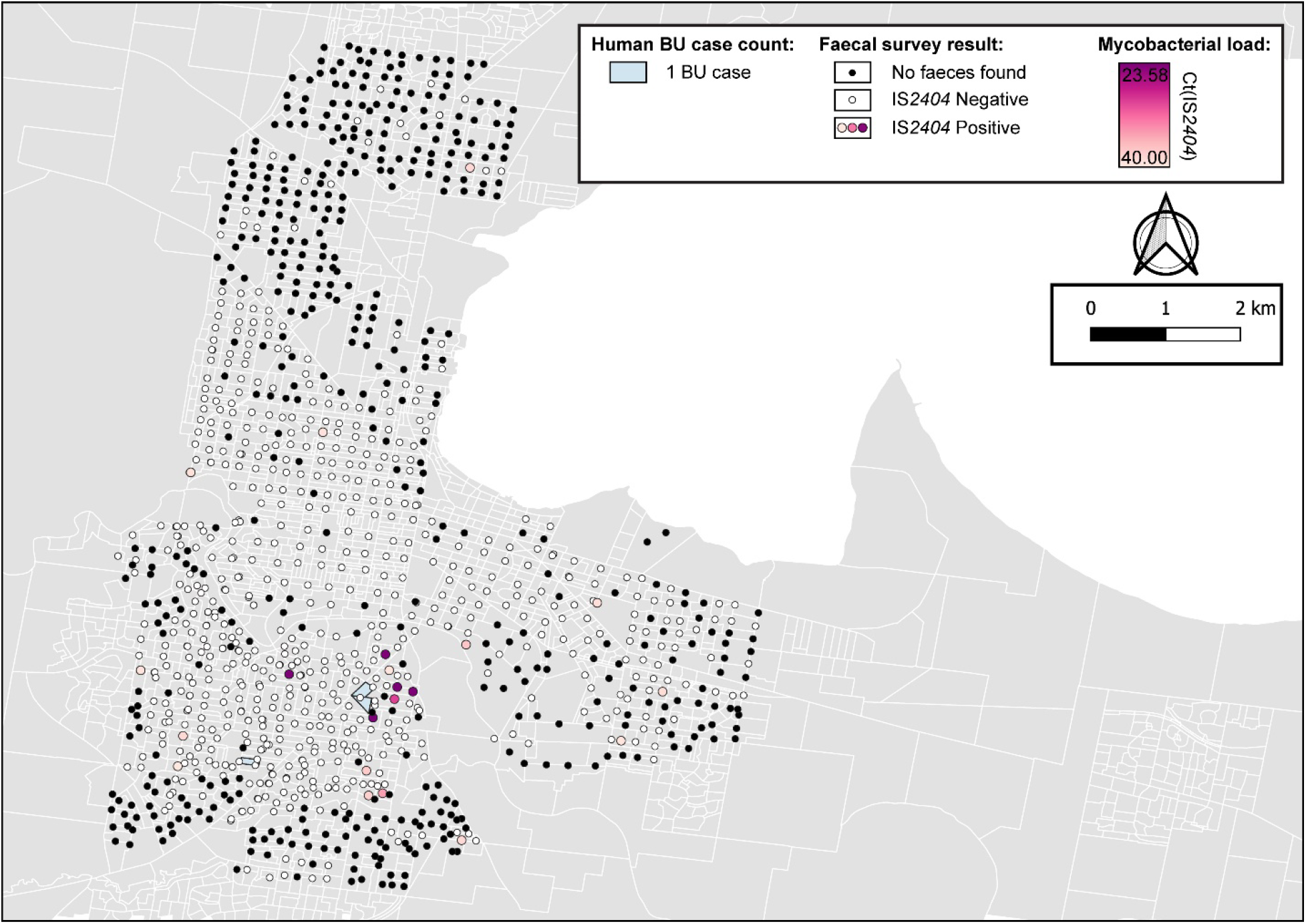
Geographical surveillance of M. ulcerans in the Geelong region. The distribution of points where possum excreta was sampled along a 200m grid pattern is presented alongside with IS2404 molecular screening results. The pink to purple color gradient visualizes inferred mycobacterial loads in analyzed excreta as estimated from IS2404 qPCR results. All BU patients notified to the DH with an inferred exposure time that overlapped with the Geelong excreta survey organized between 16/01/2020 and 28/04/2020 tabulated here by mesh block.

**Figure S6:**
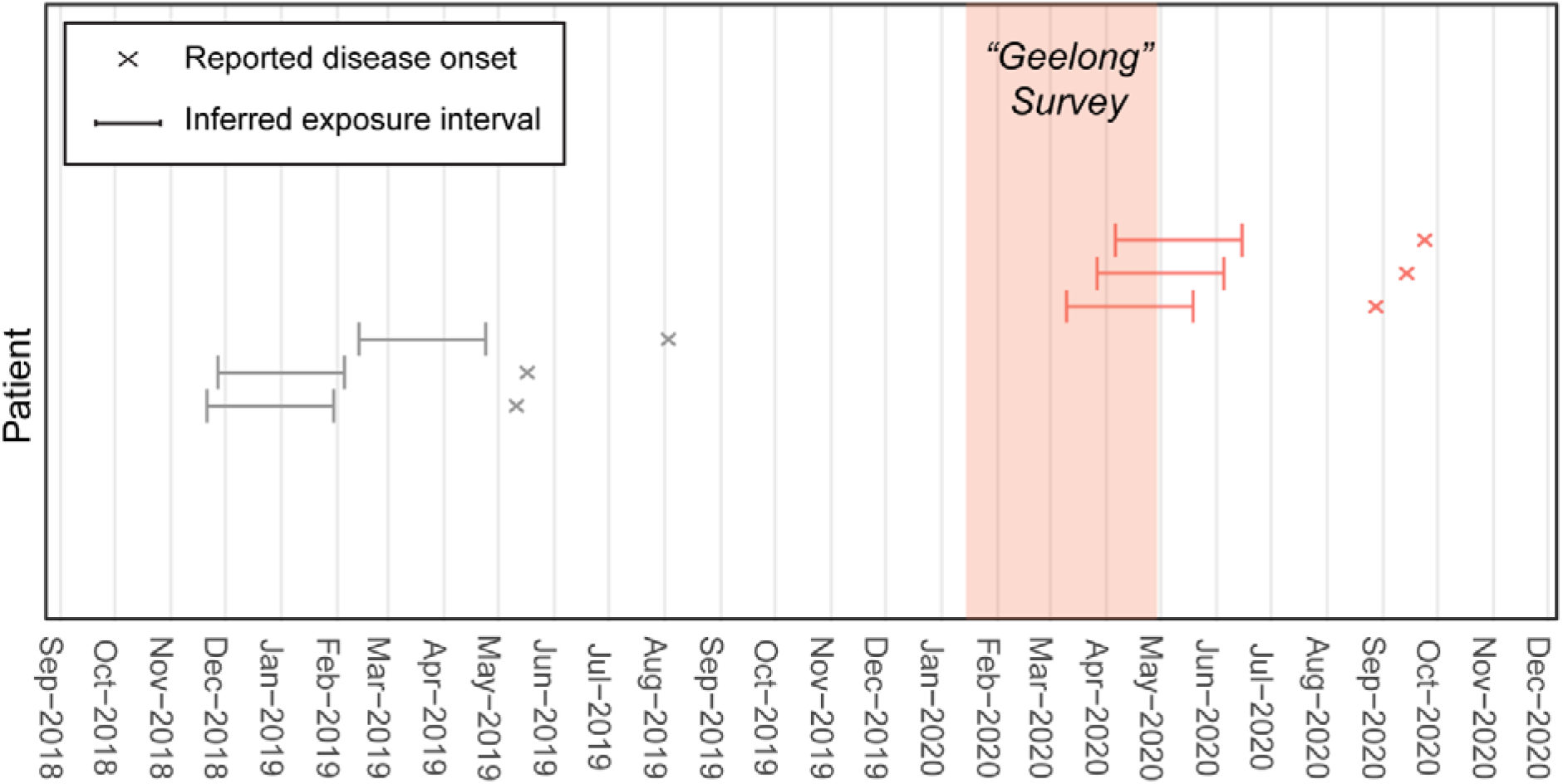
Population of BU patients with exposure intervals that aligned with the Geelong excreta possum survey organised between 16 January and 28 April 2020. The population comprises BU patients notified to the Victorian DH who resided in or visited the Geelong area and had not reported recent (<12 months) contact with any other known BU endemic areas in the state. The reported onset of disease was used to infer the exposure interval during which patients were likely infected based on the mean incubation period of BU in Victoria of 143 days (IQR 101–171) (4).

## Notes

http://www.health.vic.gov.au/infectious-diseases/local-government-areas-surveillance-report

https://github.com/abuultjens/Possum_excreta_survey_predict_human_BU

### Competing Interest Statement

The authors have declared no competing interest.

